# Mapping lineage and functional diversity in the high-risk human mammary epithelium

**DOI:** 10.64898/2026.01.12.699075

**Authors:** Matthew Waas, Bowen Zhang, Meinusha Govindarajan, Pirashaanthy Tharmapalan, Abhijith Kuttanamkuzhi, Olivia Drummond Guy, Michael Woolman, Hal K. Berman, Paul D. Waterhouse, Rama Khokha, Thomas Kislinger

## Abstract

**Background:** Breast cancer risk is shaped by the vast heterogeneity of mammary epithelial cells, comprising basal, luminal progenitor, and mature luminal populations. While transcriptional variation among these lineages has been extensively studied, protein-level features - particularly in high-risk women - remain underexplored, limiting insight into early cellular and molecular determinants of susceptibility. Moreover, little is known about how clinical covariates influence clonogenic capacity, proteomic states, and epithelial proportions, complicating the design of properly controlled human studies of early breast cancer risk.

**Results:** We combined low-input proteomics with functional clonogenic assays to profile mammary epithelial cell subpopulations from a cohort of 21 breast tissues encompassing different germline mutation backgrounds, parity status and age. We quantified over 5,555 proteins and observed marked inter-donor variation in epithelial composition, proteomic programs, and colony-forming capacity. Multivariable modeling revealed that clinical covariates - including age, parity, and germline mutation status - modulate both global proteomic architecture and lineage-specific pathway activity. Parity was associated with reduced basal cell abundance, altered luminal progenitor and mature luminal proteomes, and changes in clonogenicity. Pathway analyses identified both conserved and lineage-restricted responses to shared risk factors. Projection of clonogenic signatures onto METABRIC and TCGA tumors further linked functional programs to tumor subtypes and clinical outcomes.

**Conclusions:** This study provides the most comprehensive proteomic atlas of cell-type resolved diversity in the high-risk breast to date. By defining how clinical covariates remodel epithelial composition and molecular state, it clarifies key sources of biological variability that challenge controlled study design and offers a resource for improving mechanistic insight, risk assessment, and prevention strategies.

## Background

Breast cancer remains a leading cause of cancer-related mortality among women worldwide. Novel prevention strategies will depend on a better understanding of the earliest molecular changes in the breast epithelium.[1] Most breast cancers originate from mammary epithelial cells (MECs), which form an organized bilayer of basal/myoepithelial cells and luminal cells, which can be resolved into progenitor and mature hormone-sensing populations.[2–4] The proportion of these cell populations shifts across developmental stages such as puberty, pregnancy, and menopause, and is also influenced by inherited cancer risk factors such as germline *BRCA1/2* mutations.[5–10] Breast cancer risk factors, including age, parity, and genetic background, contribute to disease development in part by modifying epithelial cell composition and functional states.[5,6,11–14]

Single-cell RNA sequencing (scRNA-seq) has been used to great effect to decipher MEC heterogeneity, validating the canonical basal, luminal progenitor, and mature luminal compartments, while also uncovering unique subpopulations and transcriptional states within each lineage.[11–14,14–20] However, transcript abundance does not always correlate with protein abundance and proteomic validation of transcriptionally defined subtypes has been limited. Cell-state assignments can be confounded by clinical covariates, notably hormone cycle stage, parity, age, etc. Moreover, proteomic studies to date have focused on bulk-risk populations or tumor-adjacent normal tissue without cell type delineation.[21,22] Deep proteomic profiling of primary MECs from high-risk individuals without overt cancer has long been constrained by the limited number of cells obtainable from prophylactic mastectomy samples.

Recent advances in low-input proteomics enable high-depth protein quantification from scarce primary epithelial populations.[23,24] Here, we integrate low-input proteomics with flow cytometry and clonogenic assays to comprehensively profile the three major MEC subpopulation from a cohort of human breast tissues spanning a range of ages, parities, hormone-cycle phases, and germline mutation backgrounds. We quantified how these clinical covariates impact epithelial composition, protein abundance programs, and stem/progenitor-associated clonogenic potential. We further compared proteomic features with transcriptome-derived MEC signatures to evaluate concordance and covariate dependencies. Finally, we test the relevance of MEC clonogenic programs by projecting subtype-specific clonogenic signatures onto breast tumor transcriptomes from METABRIC and TCGA to assess their relationship with tumor subtype and clinical behavior.

Together, our findings provide a deep functional and proteomic portrait of human MEC heterogeneity across clinically relevant contexts. By identifying convergent and divergent effects of clinical risk factors on epithelial states **-** and by linking clonogenic MEC programs to features of breast cancers **-** this work illuminates epithelial mechanisms that may underlie risk modulation and the earliest stages of malignant transformation in the human breast. Importantly, by defining the extent and structure of inter-donor variability at the protein and functional levels, this study establishes a foundation for designing properly controlled and statistically powered future studies, enabling more precise dissection of risk-associated epithelial biology and its role in tumor initiation.

## Results

### Surveying inter-donor heterogeneity of major MEC populations and its relationship to clinical covariates

We profiled the major mammary epithelial cell (MEC) populations - basal cells (BC), luminal progenitors (LP), and mature luminal (ML) cells - of a cohort of 21 high-risk women using clonogenic assays and low-input proteomics (Figure 1A). The major MEC populations, defined as BC: CD49f^high^, EPCAM^low^; LP CD49f^high^, EPCAM^high^; ML: CD49f^low^, EPCAM^high^, were detected in each sample, though the separation and proportions varied markedly between samples (Figure 1B, C). Organizing flow cytometry profiles by clinical covariates revealed a potential visual relationship between age and MEC separation (Figure S1). While inter-donor variation in the proportion of the three epithelial cell populations (defined by EpCAM and CD49f) has been reported,[11] this phenotype has not been thoroughly examined. While hierarchical clustering of MEC proportions did not reveal clear correlations with age, parity, germline mutation, or sex hormone phase (collectively referred to as “clinical covariates”) (Figure 1C). This showed there was perhaps a more complicated relationship between covariates and epithelial proportions. We took a more systematic approach to investigate whether the heterogeneity in MEC proportions could be explained by clinical covariates using univariable and multivariable analysis (Figure 1D, Additional file 1). Compared to nulliparous women, parous women had decreased basal cell proportion (though statistically insignificant) and increased mature luminal cell proportion (Wilcoxon, P-value of 0.011); luminal progenitor proportions were independent of parity (Figure 1D). No statistically significant differences were detected across patients with different germline mutation status or hormone cycle phase (Figure 1D). Consistent with the observations from Reed *et al.*,[14] ML proportion increased while LP and BC proportions decreased with age – though the changes detected in our dataset were not statistically significant (Figure 1D). We tested whether the relationship to MEC proportions was maintained when controlling contributions of each covariate using a multivariable linear model (Additional file 1). The only significant difference was the decrease in basal cells with parity (ANCOVA of linear model, P-value 0.022). As a sensitivity check, we repeated the analysis excluding groups with few members. Results were consistent, suggesting the relationship between parity and decreased proportion of basal MECs is a robust finding (Additional file 1). The persistent inter-donor heterogeneity we observe reveals the underappreciated complexity of how clinical covariates impact remodeling of the mammary epithelium.

**Figure 1.**
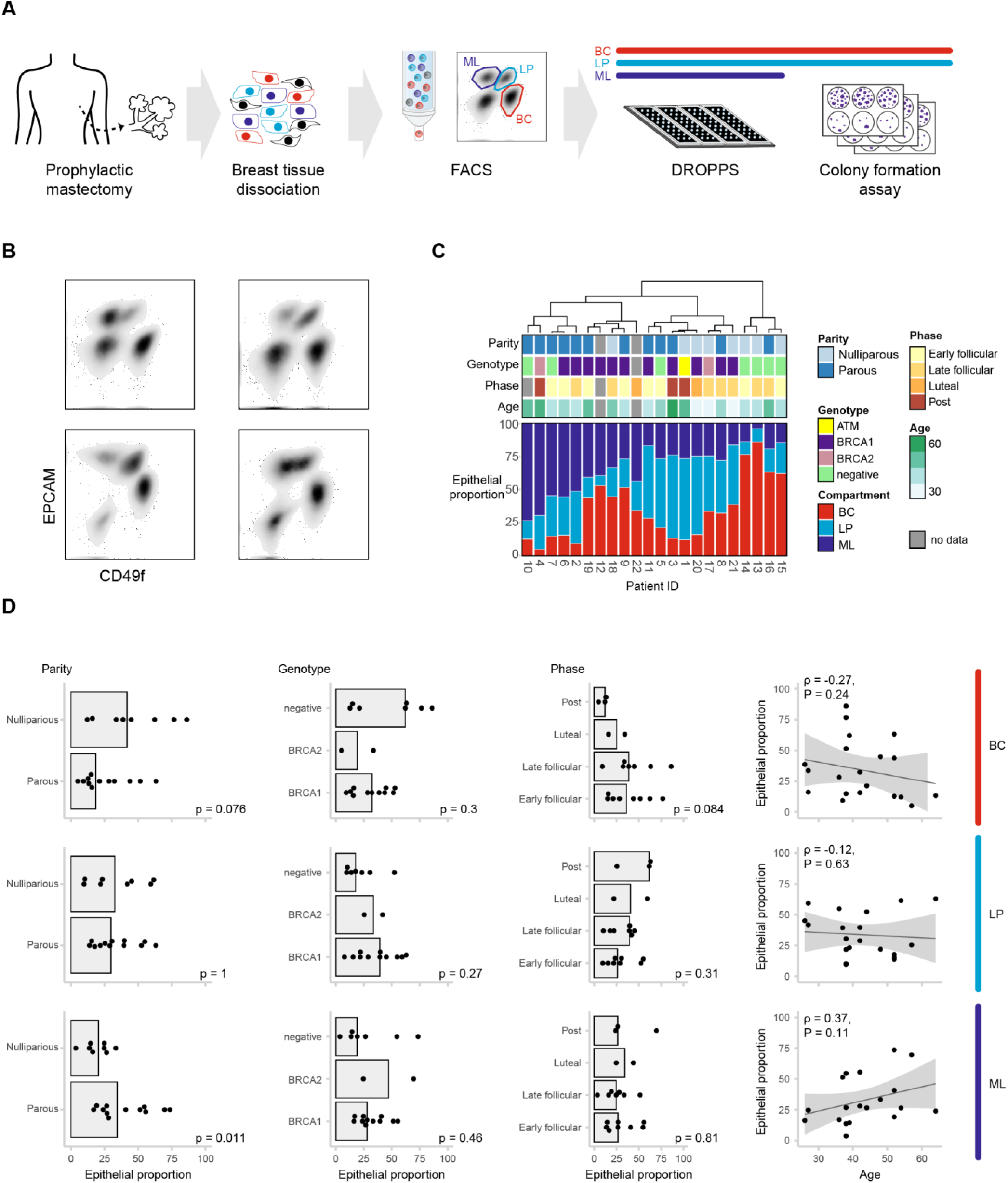
Heterogeneity of mammary epithelial cell (MEC) proportions across high-risk donors and relationship to clinical covariates. (A) Schematic of the workflow for profiling basal (BC), luminal progenitor (LP), and mature luminal (ML) MECs from prophylactic mastectomy samples using flow cytometry, clonogenic assays, and DROPPS-based low-input proteomics. (B) Example flow cytometry plots illustrating gating strategy for BC (CD49f^high^, EPCAM^low^), LP (CD49f^high^, EPCAM^high^), and ML (CD49f^low^, EPCAM^high^) populations. (C) MEC subtype proportions for all donors, hierarchically clustered and annotated by clinical covariates (age, parity, germline mutation status, sex hormone phase). (D) Univariable analysis of MEC proportions across clinical covariates (age, parity, germline mutation status, hormone phase). Wilcoxon rank sum test is used for comparison of two categories, Kruskal-Wallis is used for comparison of three or more categories. Unadjusted p-values are displayed for each test. Spearman’s correlation R and p-values are displayed with a linear model estimate where shading indicated 95% confidence interval. Summary of additional statistical tests are provided in Additional file 1.

### Clinical covariates shape MEC proteomes in patterns consistent with epidemiological risk

Recognizing that impact of clinical covariates on breast cancer risk is realized through more nuanced mechanisms than shifts in bulk proportionality alone, we developed deep proteomic portraits of high-risk breast, using DROPPS[23,24] to profile the same donor samples we analyzed by flow cytometry. Overall, we detected 5555 proteins (in 2 or more runs) with a mean depth of 4544 (range of 3395 – 5140); proteomic depth depended on MEC subtype (Figure S2A, Additional file 2). Low variance of internal controls indicated highly robust sample preparation and data acquisition (Figure S2A). Strikingly, 15 proteins detected in this cohort lack prior protein-level evidence (according to UniProt annotations), highlighting the extent to which the proteome of primary MECs remains underexplored (Additional file 2). Despite the heterogeneity observed in respective flow cytometry profiles, the proteomes clustered separately by MEC subtype (Figure 2A). We investigated the distribution of MEC lineage markers within our cohort, revealing a clear separation by cell type, but no distinguishable patterns based on clinical covariates (Figure 2B). This demonstrates the robustness of established lineage markers, whose expression is minimally affected by clinical covariates. Since MEC subtypes contributed the majority of variance, we performed PCA within each MEC subtype separately (Figure S2B). Even so, donors were distributed across PC1 and PC2 dimensions, with no clear stratification (Figure S2B). To provide protein- and pathway-level insights, we evaluated how protein expression varied across clinical covariates within each MEC subtype by multivariable linear models. For all three MEC subtypes, proteomic changes associated with *BRCA1* mutation status were significantly positively correlated with changes associated with age, consistent with observations from previous molecular profiling studies where *BRCA1* mutant breast epithelia display an accelerated aging phenotype (Figure 2C).[13] Changes associated with parity were significantly negatively correlated with age and *BRCA1* heterozygosity (Figure 2C). Strikingly, the opposing molecular effects associated with these clinical covariates align with the epidemiological associations with breast cancer risk[25]. Age and *BRCA1* mutation status are both associated with increased breast cancer risk and had positively correlated effects. While parity, associated with decreased overall risk, is negatively correlated with effects conferred by age and *BRCA1* mutation status. The relationship between hormone cycle phase and age was the most distinct across MEC subtypes, with different directionality between ML and the other two subtypes (Figure 2C). These findings support adopting MEC subtype-specific correction for phase in molecular profiling data, particularly when assessing contribution of other clinical covariates. By showing that covariates linked to risk - such as age, BRCA1 heterozygosity, and parity - drive opposing molecular programs within MEC subtypes, these MEC-lineage-resolved analyses establish a mechanistic bridge between epidemiologic observations and the cellular pathways that underlie breast cancer susceptibility.

**Figure 2.**
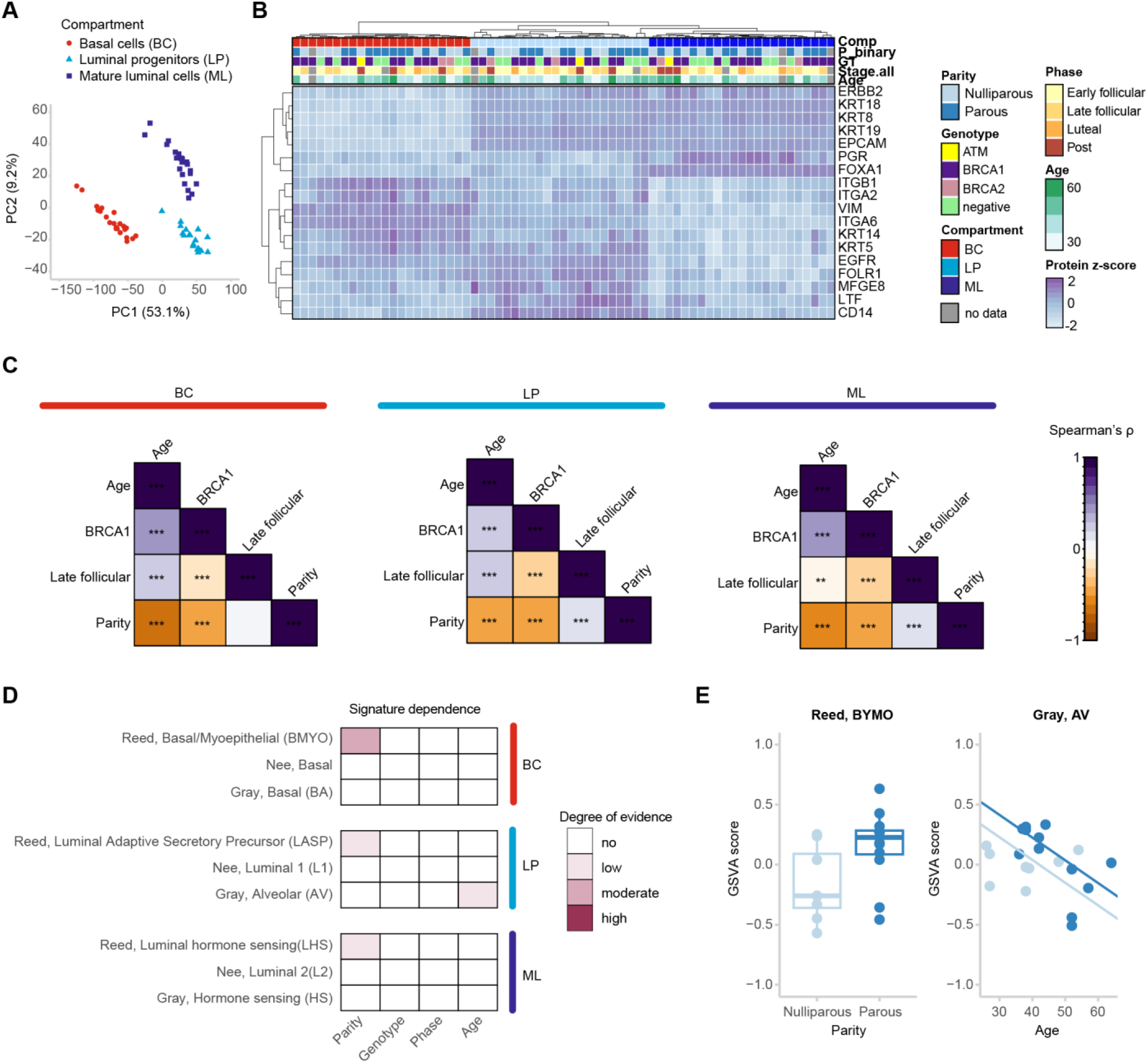
MEC proteomes cluster by subtype and show covariate-associated molecular patterns aligned with epidemiology. (A) PCA of proteomes from BC, LP, and ML cells across donors shows clustering by MEC subtype. (B) Distribution of canonical MEC markers across donors, confirming that subtype-defining proteins are minimally affected by clinical covariates. (C) Correlation matrix of covariate-associated protein expression changes for each MEC subtype. Stars indicate level of significance [no stars: p-value > 0.05, **: p-value < 0.01; ***: p-value < 0.001] (D) Strength of evidence for associations between GSVA enrichment of published subtype signatures and clinical covariates. (E) Examples of how enrichment of subtype signatures change with respect to clinical covariates. Boxplots show median and interquartile values, and whiskers show 95% confidence intervals estimates. Summary of additional statistical tests are provided in Additional file 1.

### Evaluating published MEC signatures with proteomic data across clinical variables

To assess concordance with previous studies of human MEC heterogeneity, we mapped published subtype signatures [11,12,14]-derived from scRNA-seq or CyTOF-to our proteomic data. For most signatures, we were able to detect protein-level evidence for the included genes; signatures with < 10 mapped proteins were excluded. We then calculated sample-wise enrichment for these signatures using Gene Set Variation Analysis (GSVA) on each MEC subtype independently to investigate associations between clinical covariates and signature enrichment in both univariable and multivariable models. We summarized our results based on a scale of evidence for a significant association based on the multiple models (detailed in Methods). For the primary MEC subpopulations (e.g. ML, LP, and BC), this analysis may indicate whether the signatures depend on proteins that change with respect to clinical covariates - an important consideration for cell type assignment and clustering. Reassuringly, we observed that the principal MEC subpopulation signatures were mostly independent of clinical covariates (Figure 2D). However, we did find moderate evidence that parity impacts the enrichment of the Reed *et al*. Basal/myoepithelial (BMYO) signature (Figure 2D). For comparison, we show the relationship between age and parity on Gray *et al*. alveolar luminal (AV) signature enrichment, which has lower evidence of association with clinical covariates (Figure 2E). For secondary subpopulations, we observed more evidence that the signatures differed across clinical covariates (Figure S2C). Our findings concorded with previous reports to some degree. For example, the Reed *et al.* Luminal alveolar secretory precursor 4 (LASP4) LP subtype signature - reported to be expanded in *BRCA1* mutation carriers[14] - was more enriched in *BRCA1* mutation carriers than non-carriers in our proteomic dataset (Figure S2D). While the strongest evidence for a statistically significant relationship was between LASP4 enrichment and *BRCA1* mutation status, we also observed moderate evidence that LASP4 enrichment differs across sex hormone phases and low evidence that LASP4 enrichment is impacted by age (Figure S2C, D). Overall, these analyses provided protein-level evidence of RNA-Seq defined signatures and demonstrated that the primary subpopulation signatures are mostly independent of clinical covariates. These patterns suggest that while lineage-defining programs remain constant, risk-associated exposures may act by reshaping the distribution or molecular state of rarer subpopulations, offering clues about the earliest epithelial changes that could predispose to malignant progression.

### Pathway analysis reveals convergent and divergent responses to clinical covariates among MEC subtypes

To more precisely define how clinical covariates impact MEC biology beyond shifts in cell type frequency, we performed Gene Set Enrichment Analysis (GSEA) independently within each primary MEC subtype (using Hallmarks, GO Biological Processes, and KEGG Legacy signatures). Many pathways demonstrated significant associations which are consistent with expectation, for example, estrogen signaling was dependent on sex hormone phase (Additional file 3). Interestingly, while many pathways were specific to a single covariate or subtype, there is a subset of pathways that emerged as significantly associated across multiple covariates and subtypes; the most common pathways (> 6 significant associations) fell into the broad categories of metabolism, immune signaling, and stress response (Figure 3A-B). These represent the pathways that are most susceptible to influence by clinical covariates. Notably, one pathway with significant associations across multiple covariates for each primary subpopulation was “mitochondrial gene expression” (Figure 3A). This was notable as many scRNA-Seq pipelines use mitochondrial read content as a criterion for filtering or as a feature for normalization. If mitochondrial content or transcriptional regulation are determinants of plasticity for MEC subtypes, then standard preprocessing may inadvertently obscure or confound true biological signal by treating mitochondrial read content differences primarily as technical artifacts.

**Figure 3.**
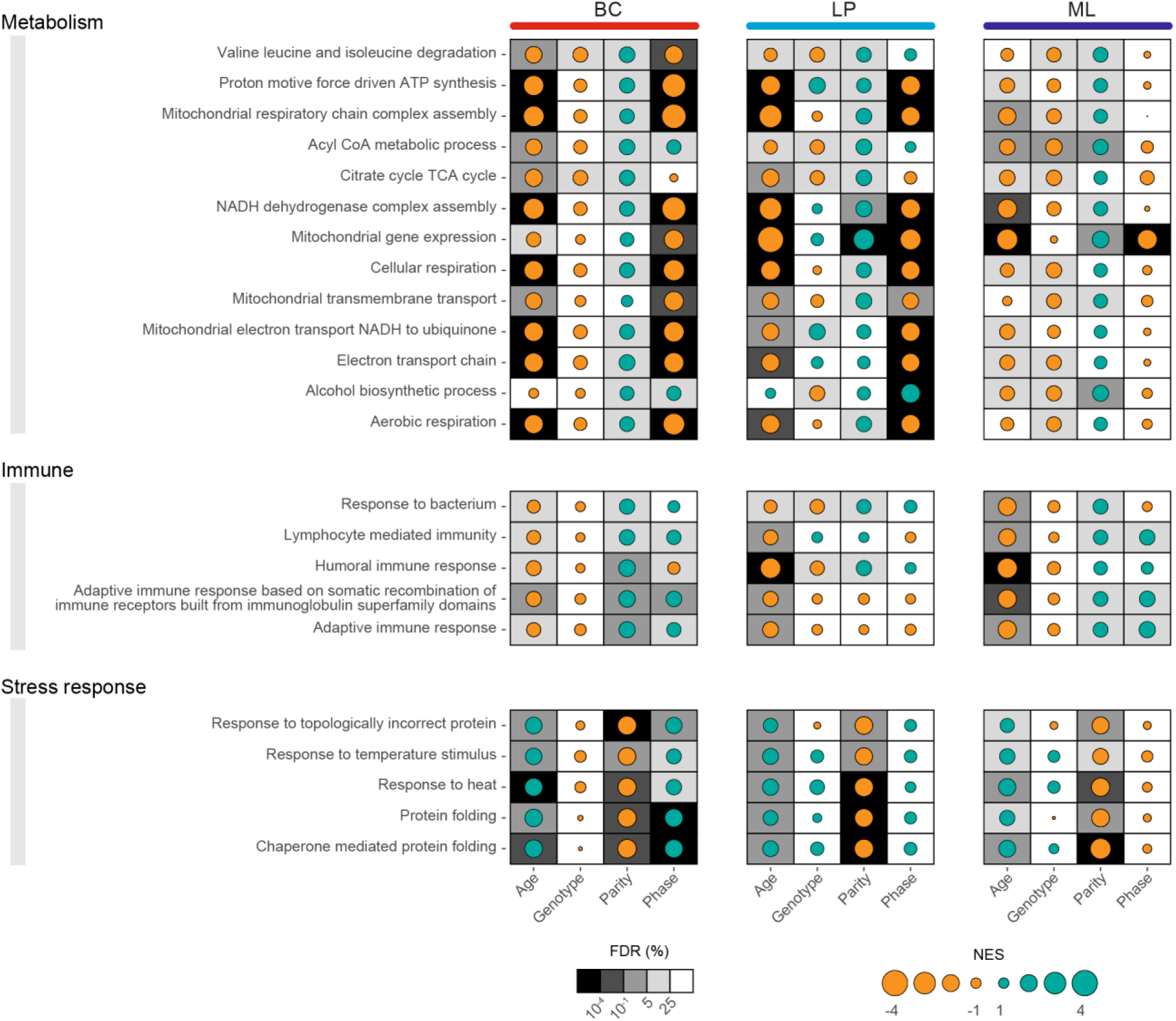
Convergent and divergent pathway responses to clinical covariates across MEC subtypes. Dotplot of pathways which were significantly associated using GSEA with clinical covariates in MEC subtypes. NES directionality is represented by color and the magnitude by the size of dots. The significance of the enrichment is depicted as the shading in the background. All pathways with >6 significant values were shown. All GSEA outputs are provided in Additional file 3. FDR: Benjamini-Hochberg adjusted p-values from GSEA.

To investigate inter-donor heterogeneity of MECs in the context of cancer-associate pathways, we calculated sample-wise enrichment for Hallmark Signatures using GSVA (Figure S3). Some Hallmark terms had clear relationships with MEC subtype. For example, gene sets describing “apical junction” and “myogenesis” pathways were strongly enriched in basal cells, consistent with their *in situ* spatial localization and expression of myoepithelial contraction-related proteins. Aligning with our previous reports,[26,27] pathways such as “oxidative phosphorylation” and “DNA repair” were highly enriched in luminal progenitors. Next, we evaluated the relationship between each Hallmark term and clinical covariate in a subtype specific manner, (Figure S3) scoring evidence for association using a similar methodology as the previous signature enrichment analysis (Figure 2D, S2C) (detailed in Methods). This analysis revealed both shared and diverging effects of clinical covariates on MEC subtypes. Age emerged as the covariate with the strongest evidence of impact on Hallmark Signatures. Notably, the signatures for both “inflammatory response”, and specifically “IL6-JAK/STAT3 signaling” showed high and moderate evidence, respectively, for association with sex hormone phase in each MEC subtype (Figure S3). The interaction between IL6-JAK/STAT3 and hormone signaling has been previously reported in both culture and *in vivo* systems.[28–32] Broadly, these data uncover shared and lineage-specific pathways through which clinical covariates remodel the mammary epithelium, providing a more granular framework for understanding how inter-individual differences may modulate breast cancer risk.

### Proteomic correlates of MEC clonogenic content are lineage-specific and related to clinical presentation of breast cancer

Stem and progenitor cells within the mammary epithelium are the putative origin of aggressive breast cancer subtypes. Clonogenicity, or colony forming potential, provides a functional readout of stem/progenitor content within the mammary epithelium. As our results revealed that clinical covariates can impact mammary epithelial proportions and properties, we assessed whether they influence clonogenic content of BC and LP populations using colony forming assays on cells sorted from the same samples analyzed by flow cytometry and proteomics (example images in Figure 4A). First, we evaluated whether clonogenicity was coordinated across MEC subpopulations by plotting the colony counts from LP and BC populations against one another and annotating by clinical covariates (Figure S4A). The weak correlation and absence of clear covariate-driven patterns suggested that clonogenic content were independently regulated in BC and LP (Figure S4A). We next modeled the effects of clinical covariates on clonogenicity using univariable and multivariable approaches, evaluating contributions to both colony count and colony size. In univariable analyses, BC colony counts decreased with parity, whereas LP colony counts increased during the late follicular phase (Additional file 1, Figure 4B, S4B). These covariate effects were subpopulation specific. Parity did not affect LP colony counts, and sex hormone phase had no detectable effect on BC colony counts. Colony size and colony number were also not always concordant. For example, LP colonies in the late follicular phase were both larger and more numerous, whereas parity was associated with fewer but larger BC colonies (Figure 4B, S4B). Together, these findings show that stem/progenitor activity in the mammary epithelium is shaped by covariate-specific and lineage-specific mechanisms, underscoring how common clinical factors may differentially modulate the pools of cells most relevant to breast cancer initiation. Importantly, these results demonstrate that if clonogenic content is to be used to evaluate differences across one biological variable such as genotype, careful matching of the other covariates is essential for proper interpretation.

**Figure 4.**
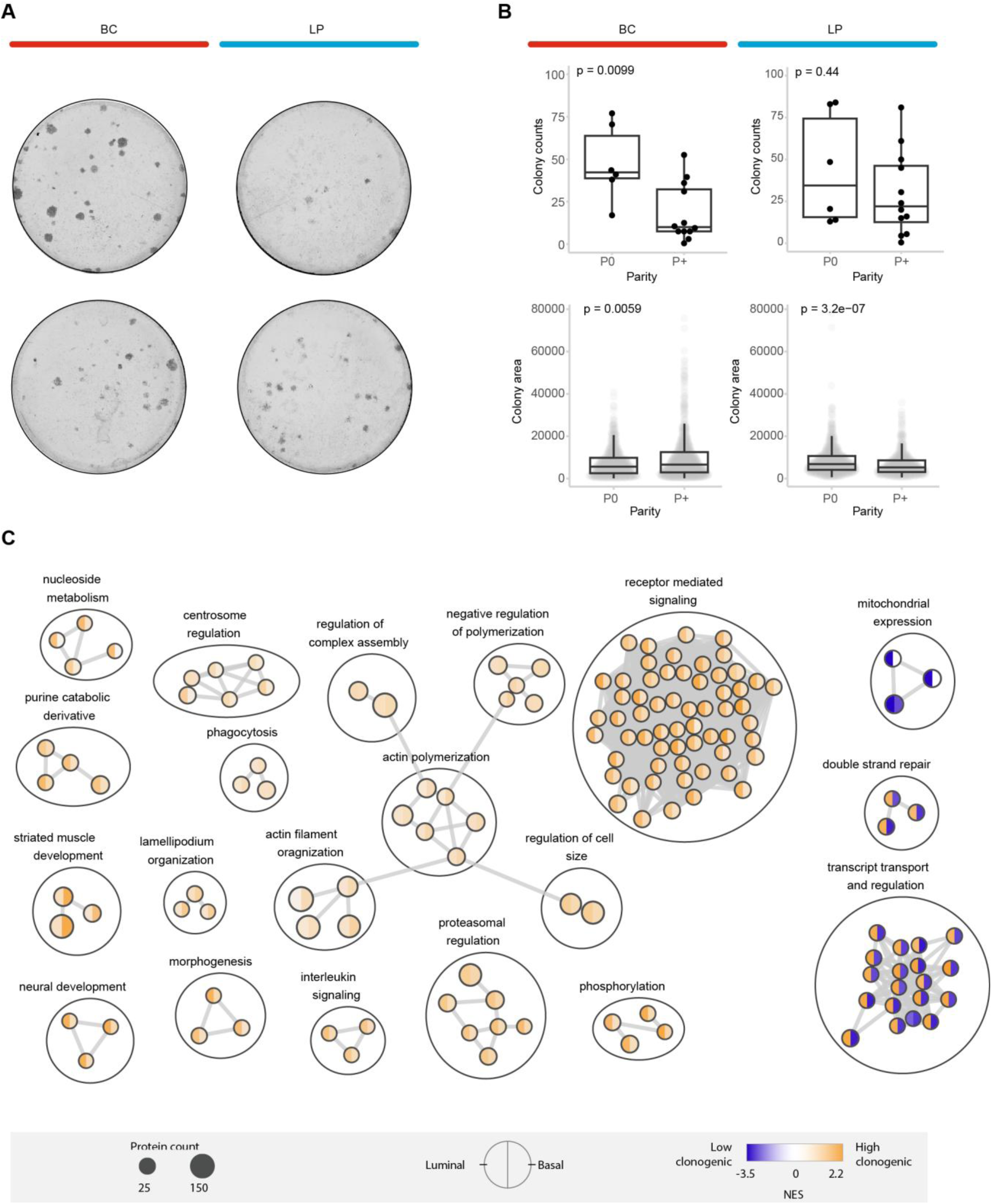
Lineage-specific clonogenic capacity reflects MEC functional states. (A) Example images of BC and LP colonies. (B) Distribution of LP and BC colony counts and sizes across parity groups. Unadjusted p-values from Wilcoxon rank sum test are displayed. Boxplots show median and interquartile values, and whiskers show 95% confidence intervals estimates. Summary of additional statistical tests are provided in Additional file 1. (C) Cytoscape enrichment map for significantly enriched pathways associated with clonogenic content from GSEA. The NES direction indicated by color and the magnitude of the enrichment is represented as the shading. All GSEA outputs are provided in Additional file 3.

To identify molecular correlates of clonogenic capacity, we incorporated clonogenic content (median dichotomized colony counts) as a categorical variable into the multivariable models for LP and BC proteomes. We performed GSEA to investigate pathways associated with LP and BC clonogenicity (Figure 4C, all GSEA results in Additional file 3). These analyses highlighted pathways related to receptor-mediated signaling, developmental and morphogenetic programs, nucleotide metabolism, and actin remodeling (Figure 4C). Despite donors rarely exhibiting high colony counts in both LP and BC (Figure S4A), the molecular pathways associated with high clonogenicity were strikingly similar across subtypes. This indicates that these clonogenic programs can be independently engaged in each MEC subtype and are not explained by donor-driven differences alone. Although most enriched pathways overlapped between subtypes, some differed in magnitude, and a subset - particularly DNA repair and transcript-transport/regulation pathways - were anti-correlated between BC and LP (Figure 4C). Extracting the top 100 proteins with high estimated contributions from each MEC subtype resulted in lineage-resolved clonogenic content signatures with an overlap of only 4 proteins: TTC4, C3, DPYD, TRAF1.

Finally, we reasoned that if the clonogenic programs in high-risk MECs are relevant to tumorigenesis, they may be related to different clinical presentations of breast cancer. To test this hypothesis, we projected the LP and BC clonogenic signatures onto bulk tumor transcriptomes from METABRIC and TCGA (Figure 5A, B). Both BC and LP clonogenic signatures were higher in triple-negative than in hormone receptor-expressing breast cancers (Figure 5A), supporting the hypothesis that these cancers originate from mammary stem/progenitor cells. Interestingly, while the BC clonogenic signature was increased in both classes of cancer compared to non-cancer samples, the LP clonogenic signature was highest in non-cancer (Figure 5A). The transformation from luminal progenitors to tumors is hypothesized to involve an intermediate state, termed basal-luminal cells.[11,33], which are characterized by a loss of luminal lineage fidelity. One interpretation of the observation that luminal progenitor clonogenic signatures are higher in non-cancer than cancer cells, is that the transformed tumor cells exhibit a more basal-like proliferative program. We further investigated our scores, by mapping the enrichment scores across PAM50 (and claudin-low) transcriptional subtypes (Figure 5B). The results from TCGA and METABRIC were mostly concordant with the lowest scores for BC- and LP-clonogenic signatures belonging to Luminal A and Luminal B subtypes - cancers that are thought to arise from more differentiated MEC populations (*e.g*. ML cells) (Figure 5B), which do not demonstrate colony forming capacity under standard conditions. In the METABRIC cohort, the BC- and LP-clonogenic score distributions were highly similar, with scores highest in the claudin-low subtype, followed by basal-like and normal-like cancers (Figure 5B). In the TCGA cohort, the BC- and LP-clonogenic score distributions were slightly more divergent. The BC-clonogenic signature scored highest in claudin-low and Her2 subtypes, followed by basal-like (Figure 5B). The LP-clonogenic signature scored highest in claudin-low, followed by normal-like (Figure 5B). As discussed elsewhere,[34] there were different inclusion and exclusion criteria for each cohort, which may have impacted the observed frequency and characteristics of each subtype. Altogether, these data reveal that clonogenic programs in high-risk MECs are both lineage-specific and clinically meaningful. Their stratification across breast cancer subtypes - and enrichment in the most clinically aggressive breast cancers - suggest that functional epithelial traits observable in non-malignant tissue may portend patterns of tumor progression. These results highlight the potential value of integrating functional assays with proteomics when evaluating early determinants of breast cancer risk.

**Figure 5.**
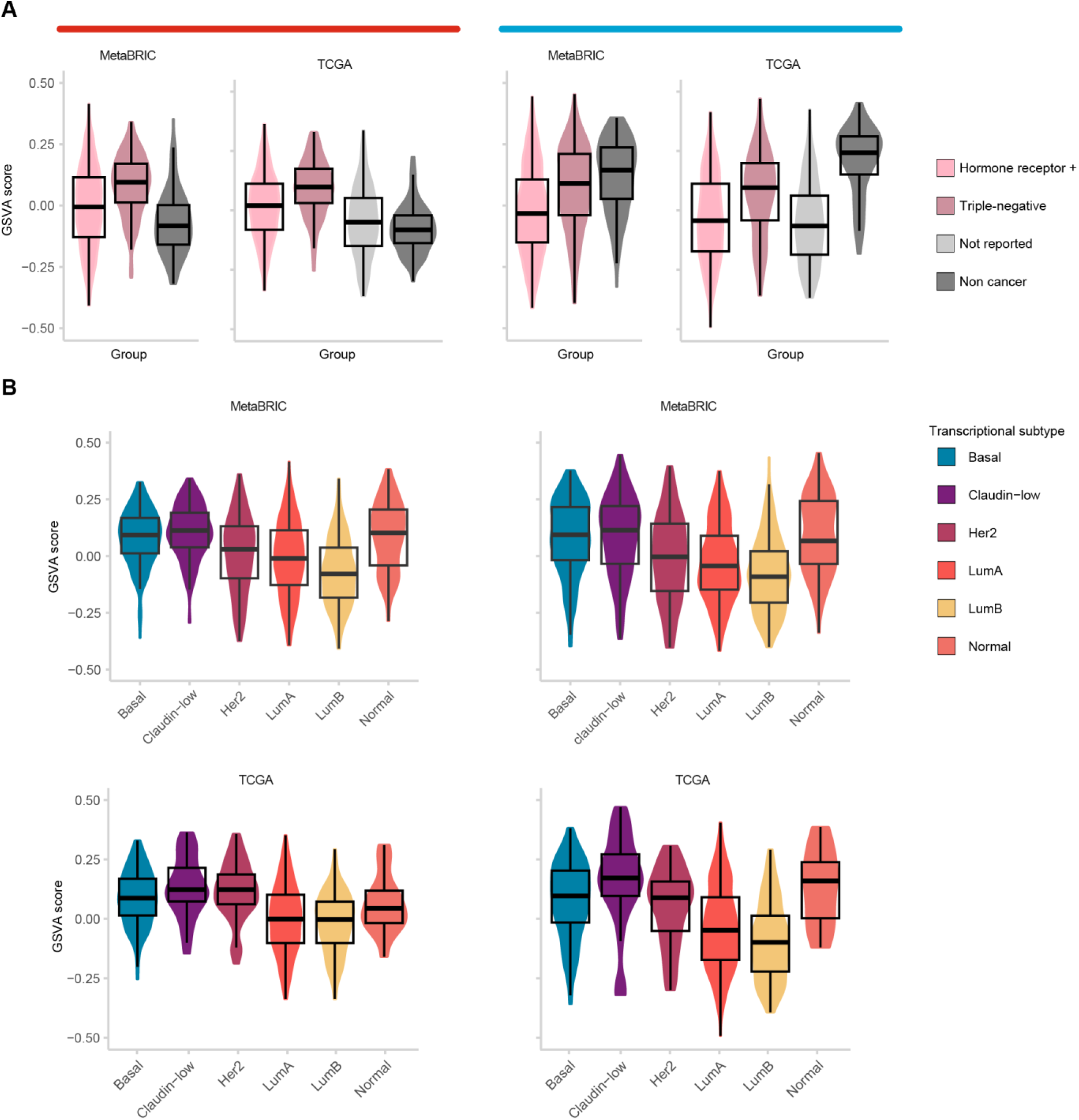
Lineage-resolved MEC clonogenic programs map onto breast tumor phenotypes. (A) Distribution of GSVA enrichment scores for BC- and LP-clonogenic scores across triple-negative cancer, hormone-expressing cancer, and non-cancer samples in METABRIC and TCGA cohorts. (B) Distribution of GSVA enrichment scores for BC- and LP-clonogenic scores across transcriptional subtypes of breast cancers in METABRIC and TCGA cohorts.

## Discussion

Our study provides a comprehensive proteomic and functional characterization of mammary epithelial cell (MEC) subtypes from high-risk women, revealing how clinical covariates - including age, parity, sex hormone phase, and germline mutation status - shape epithelial composition, proteomic states, and clonogenic potential. At the level of epithelial composition, several of our observations provide important context for earlier literature. We found that parity decreased BC proportionality, contrasting with Reed et al.[14], who reported an increase. Methodological differences in estimating epithelial proportionality may contribute to this discrepancy, similar to inconsistencies noted in studies examining LP expansion in BRCA1 mutation carrier.[6,17] Conflicting associations between basal proportionality and age further suggest that some previously reported signatures may capture unmodeled covariate confounders. These inconsistencies underscore the challenges of quantifying epithelial composition in primary human tissue and highlight the need for standardized analytical frameworks and orthogonal validation and enumeration strategies.

Our proteomic findings shed light on how each clinical covariate impacts MEC pathway engagement. Many of the proteomic differences we observed aligned with established epidemiological patterns as to how covariates impact cancer risk. LP cells showed the most covariate-responsive pathway engagement, consistent with their proposed role as the cell-of-origin for aggressive breast cancers such as basal-like or HER2-enriched disease. These concordances reinforce the biological validity of the proteomic phenotypes detected and indicate that population-level risk modifiers leave measurable molecular imprints on MEC states. Despite the heterogeneity of exposures examined, many perturbed pathways overlapped across lineages, suggesting that physiological and genetic risk factors converge on a conserved set of cellular programs that govern epithelial plasticity and perhaps vulnerability to malignant transformation. These shared pathways - including immunomodulatory, metabolic, and hormone-responsive networks - may represent core circuits through which the mammary epithelium responds to environmental or endogenous perturbations. This epithelial plasticity likely contributes to how different lineages or functional states become, thereby shaping breast cancer risk. Further questions remain, such as the extent of interactions between *BRCA1* mutation status, hormone receptor-based signaling, and inflammatory signaling [28,29,31,32,35,36]. Careful mechanistic investigation of these interacting influences will help elucidate how epidemiologic risk factors translate into lineage-specific vulnerabilities.

Clonogenic analyses demonstrated that MEC functional capacity is both lineage-specific and covariate-modulated. BC and LP clonogenicity were only weakly correlated across donors, indicating largely independent regulation of progenitor activity within each lineage. Univariable and multivariable models revealed distinct covariate effects: parity decreased BC clonogenicity without affecting LP clonogenicity, whereas hormonal phase increased LP colony counts and size but had no detectable effect on BC. These findings reinforce that clinical exposures shape lineage function through subtype-specific mechanisms rather than through uniform shifts. Pathway analyses showed that despite minimal donor-level concordance in clonogenic phenotypes, BC and LP clonogenicity were associated with strikingly similar biological programs - receptor-mediated signaling, morphogenesis, metabolism, and actin remodeling - suggesting that shared functional modules can be independently activated across lineages.

By integrating low-input quantitative proteomics with lineage-resolved clonogenic assays, we capture features of epithelial heterogeneity that complement and extend transcriptome-based studies. Protein-level measurements offer direct insight into effector molecules, pathway activation, and post-transcriptional regulation, providing a more mechanistic readout of cellular state. Projection of BC and LP clonogenic signatures onto METABRIC and TCGA tumors further demonstrated that functional programs active in high-risk MECs correspond to tumor subtype and clinical presentation. Both BC and LP clonogenic signatures were enriched in triple-negative and claudin-low tumors - subtypes frequently hypothesized to originate from basal-like cells or aberrantly differentiated luminal progenitors - while luminal tumors exhibited lower scores, consistent with more differentiated MEC origins. These observations suggest that normal lineage-specific functional states are maintained, co-opted, or reconfigured during tumorigenesis and serve as a biological bridge between early risk-associated programs and established cancer phenotypes. Integration of clonogenic signatures with tumor transcriptomes further contextualized these programs. While BC clonogenic signatures tended to increase from non-cancer to cancer tissue, LP clonogenic signatures were highest in non-cancer samples, potentially reflecting loss of luminal lineage fidelity during transformation - a concept consistent with the proposed basal–luminal intermediate state.[11,33] This inversion between normal and malignant tissues underscores the importance of characterizing lineage states prior to transformation and suggests that susceptibility may relate not only to abundance of particular lineages but also to the stability of their progenitor programs.

Despite these insights, key challenges remain in studying primary human MECs. Inter-donor variability, covariate-driven effects, and limited access to high-quality tissue constrain statistical power and obscure causal relationships. Controlled study designs with systematic covariate stratification, orthogonal cell-type enumeration, and protein-level validation will be critical for refining mechanistic links between epithelial state and breast cancer risk. While our dataset resolves the proteomes of the major MEC lineages (BC, LP, ML), future work combining careful functional evaluation with higher-resolution proteomic platforms - such as flow-sorted subpopulations or single-cell proteomics - will be needed to define whether observed proteomic differences reflect proportional shifts, altered protein expression within stable populations, or the emergence of covariate-specific intermediates (*e.g.*, basal–luminal hybrids[11]).

Overall, this study provides a lineage-resolved proteomic resource that illuminates both conserved and lineage-specific responses to physiological and genetic risk factors. By integrating protein-level measurements with functional assays and linking these programs to tumor phenotypes, we establish a framework for understanding how epithelial plasticity and lineage-specific functional states contribute to breast cancer susceptibility. This resource lays a protein-level foundation for future mechanistic studies and opens avenues for improved risk stratification and prevention strategies.

## Conclusions

This study defines the proteomic architecture of the major mammary epithelial lineages in women at elevated risk for breast cancer and demonstrates how age, parity, and germline mutation status shape epithelial composition, pathway engagement, and functional potential. By integrating low-input proteomics with clonogenic assays and computational mapping of normal signatures onto breast tumors, we identify lineage-resolved programs that link clinical covariates to molecular states associated with transformation risk.

Across MEC subtypes, diverse epidemiologic and genetic factors converge on a shared set of pathways, underscoring the intrinsic epithelial plasticity that governs how the mammary gland adapts to physiological and hormonal cues. These patterns were most pronounced in luminal progenitors, consistent with their proposed role as the cell-of-origin for multiple aggressive tumor subtypes. Our findings also highlight areas of discordance with prior proportionality studies and reveal potential confounding interactions - such as the interplay between BRCA1 heterozygosity and ovarian hormone signaling - that warrant further mechanistic investigation.

While our work resolves proteomes at the level of major epithelial lineages, finer-resolution strategies such as single-cell or sorted-subpopulation proteomics will be needed to distinguish whether covariate-associated pathway engagement reflects intrinsic responses of all cells within a lineage or the emergence of previously unrecognized subpopulations. Together, this study establishes a protein-level framework for interpreting risk-associated epithelial states, complements existing transcriptomic atlases, and provides a foundation for future mechanistic and preventive research in breast cancer biology.

## Methods

### Donor-tissue details

All human tissues were acquired with patient consent and approval from the Research Ethics Board of University Health Network (Toronto, ON). Menstrual stage (premenopausal, early and late follicular, and luteal) was determined by histological examination of breast specimens stained with hematoxylin & eosin from at minimum 5 tissue blocks per patient. Staging was confirmed by pathologist (H. Berman). Breast specimens were collected from true and contralateral prophylactic mastectomies along with reduction mammoplasties, with relevant patient information presented in Table S1. To assign and confirm presence of specific pathogenic germline mutations, all samples were screened with a 45-gene germline mutation panel. Library preparation used 200 ng of DNA as a template, employing the xGen Library Prep Kit (IDT, Coralville, IA, USA). Individual libraries were pooled before undergoing target enrichment using custom-designed probes specifically capturing the coding exons of the HerCaP gene panel. Following enrichment, libraries were sequenced at 2x150 bp on an Illumina MiSeq Sequencer. The resulting sequence reads were aligned to the human reference genome, and variants were called using Sentieon software (Sentieon, Santa Clara, CA, USA). Called variants were classified based on the American College of Medical Genetics (ACMG) guidelines for determining pathogenic/likely pathogenic variants using VarSeq software (GoldenHelix, Bozeman, MT, USA).

### Human mammary single-cell suspensions

Breast specimens were minced, then enzymatically dissociated in a DMEM/F-12 based dissociation media (Gibco, #11330-32) containing 15 mM HEPES, 2% BSA, 1% penicillin-streptomycin, 5 µg/mL insulin, 300 U/mL collagenase (Sigma, C9891) and 100 U/mL hyaluronidase (Sigma, H3506). with gentle shaking at 37°C for 16-18 hours. The epithelia-enriched fraction, termed pellet A, was harvested by centrifugation at 80 g (30 sec) and then viably cryopreserved, as described previously. To generate single cell suspensions in preparation for FACS, viably cryopreserved pellet A vials were thawed and dissociated following standard methodology [PMID: 34031589]. Briefly, samples were triturated for 2 min in 0.25% trypsin-EDTA (Stem Cell Technologies, 07901) followed by trituration for 2 min in 5 U/mL Dispase (Stem Cell Technologies, 07913) and 50 µg/mL DNase I (Sigma, D4513). Cells were washed in with HBBS + 2% FBS following each enzymatic dissociation step and passed through a 40 µm strainer after treatment with Dispase.

### Fluorescent activated cell sorting

For FACS staining, anti-CD326-AF647 (BioLegend 324212 ; clone9C4; 1:50 dilution), anti-CD49f-FITC (BioLegend 313606; clone GoH3; 1:100), anti-CD45-PE/Cy7 (BioLegend 304016; clone HI30; 1:200), anti-CD31-PE/Cy7 (BioLegend 303118; clone WM59; 1:50), were used. Dead cells were excluded following doublet exclusion using DAPI (Sigma, D9542, 1:10000). Lineage-positive cells were defined as CD31/CD45^+^. Mammary cell subpopulations were defined as: total basal (Lin^−^EpCAM^lo-med^CD49f^hi^), luminal progenitor (Lin^−^EpCAM^hi^CD49f^hi^), mature luminal (Lin^−^EpCAM^hi^CD49f^lo^) as previously described. Cell sorting was performed on a BD FACS ARIA Fusion with FACSDiva (v.8.0.1 and v.6.1.3). Flow data analysis and plotting was performed using FlowJo (v.10.8.1).

### Human mammary colony-forming cell assay

In total, 400 cells of the specified FACS-purified epithelial populations were seeded together with 20,000 irradiated NIH 3T3 mouse fibroblasts per well in six-well plates pre-coated with collagen I (1:43 dilution of 3mg/ml rat tail collagen I in 1X PBS, Gibco A1048301) for 1 h at 37°C, StemCell Technologies). Cells were cultured for 10 days at 5% O_2_ in EpiCult-B Human medium (STEMCELL Technologies, 05601) with The EpiCult-B Human media prepared per manufacturer instructions and supplemented with 5% FBS was used to seed the FACS purified cells for 24 hours. The seeding media was replaced with serum-free complete Epicult-B Human media for the remaining 9 days. Colonies were stained with Wright-Giesma and each well per plate was scanned using the Biotek Cytation 5 Imaging Multimodal Reader (Agilent). From scanned whole well images, colony count and size was quantified through a neural network-based classifier trained on Biodock using past human CFC images. Colony classification accuracy was manually verified.

### Proteomic sample preparation

Two processing replicates of 500 cells were deposited into wells of Teflon-coated slides (Tekdon, template available upon request) by FACS and processed using DROPPS. The sheath accompanying the cells was allowed to evaporate at room temperature in a biosafety cabinet and then slides were stored at -80 °C until further processing. For lysis and protein reduction, 2.5 μL of buffer consisting of 10% (v/v) Invitrosol (ThermoFisher, MS10007), 15% (v/v) acetonitrile, 0.06% (w/v) n-dodecyl-β-D-maltoside (Sigma, 850520P), 5 mM Tris(2-carboxyethyl) phosphine, and 100 mM ammonium bicarbonate in HPLC grade water. 10% (v/v) of INV2 was included in lysis buffer. Slides were placed in a 37 °C humidity chamber and allowed to incubate for 30 min. For alkylation and protein digestion, 1 μL of buffer containing 35 mM iodoacetamide and 10 pg of Trypsin/Lys-C (Promega, V5072) was added to each sample. Samples were allowed to digest for 2 h in a 37 °C humidity chamber. Digestion was quenched by bringing samples to 0.1% (v/v) formic acid with 8 μL of 0.14% (v/v) formic acid in 37 °C HPLC grade water and then samples were transferred to 96-well plate. The wells were washed with an additional 8 μL of 0.1% (v/v) formic acid and then the plate was transferred to -20 °C freezer. After allowing samples to thaw, iRT peptides were spiked in and plate was spun at 2000 g for 5 min.

### Mass spectrometry data acquisition

LC-MS/MS analysis was performed on an Orbitrap Eclipse MS (ThermoFisher) coupled to Neo Vanquish (ThermoFisher). Peptides were washed on pre-column (Acclaim™ PepMap™ 100 C18, ThermoFisher) with 60 μL of mobile phase A (0.1% FA in HPLC grade water) at 10 μL/min separated using a 200 nL.min flow rate over a 50 cm μPAC™ Neo HPLC Columns (COL-NANO050NEOB, ThermoFisher) ramping mobile phase B (0.1% FA , 80% HPLC grade acetonitrile in HPLC grade water) from 0% to 30% in 136 min, 30% to 70% in 20 min interfaced online using an EASY-Spray™ source (ThermoFisher). The Orbitrap Eclipse MS was operated in data dependent acquisition mode using two 1.5 s cycles at different FAIMS settings (CVs of -45 and -60) with a full MS resolution of 240,000 with a full scan range of 375- 1050 *m/z* with RF Lens at 60%, full MS AGC at 250%, and maximum inject time at 50 ms. MS/MS scans were recorded in the ion trap with 0.6 Th isolation window, 20 ms maximum injection time, with a scan range of 200-1400 *m/z* using Rapid scan rate. Ions for MS/MS were selected using monoisotopic peak detection, intensity threshold of 1,000, positive charge states of 2-5, 20 s dynamic exclusion, and then fragmented using HCD with 27.5% NCE.

### Mass spectrometry raw data analysis

Raw files were analyzed using FragPipe (v.22.0) using MSFragger[37,38] (v.4.1) to search against a human proteome (Uniprot, 43,392 sequences – canonical plus isoforms, accessed 2023-02-08). Default settings for LFQ workflow[39,40] were used using IonQuant[41] v.1.9.8 and Philosopher[42] v.5.0.0 with the following modifications: Precursor and fragment mass tolerance were specified at -50 to 50 ppm and 250 ppm, respectively; parameter optimization was disabled; MaxLFQ min ions was set to 1; MBR RT tolerance was set to 1 min, and MBR top runs was set to 10, no normalization was applied.

### Mass spectrometry statistical analysis

All analysis was performed using R programming language (v.4.2.2) with Tidyverse pacakge (tidyverse_1.3.2) unless otherwise specified. All correlation estimates and p-values were calculated using the “cor.test” function. The “MaxLFQ Intensity” columns were extracted from the “combined_protein.tsv” output file from FragPipe and mean log2(intensities) were calculated from the two processing replicates and used for all subsequent analysis (Additional file 2). Proteins were filtered out using a list of common contaminants. Principal component analysis was performed on log2-transformed intensities imputing missing values to 0.

#### Within MEC subtype analysis

First proteins were filtered for presence in > 66.7% of donors for each subtype and subsequently imputed with random forest algorithm using the MissForest package (missForest_1.5). These are referred to as imputed, intra-MEC data matrices.

#### Between MEC subtype analysis

Data matrix from intra-MEC comparisons were stitched back together, and values from original matrix were filled in for all proteins detected in >50% of samples. Data were then subjected to Perseus-style[43] lower-tail imputation with downshift of 2 standard deviations and width of 0.4 standard deviations.

Heatmap generation was clustered using Pretty Heatmap package (pheatmap_1.0.12) using correlation as a distance metric for proteins, Euclidean distance for samples, and complete linkage for canonical markers. The GSVA[44] package (GSVA_1.46.0) was used to calculated enrichment of previously published signatures[11,12,14,33] on the intra-MEC data matrices.

For many analyses, we evaluated the relationship between clinical covariates and output (i.e. epithelial proportion, GSVA enrichment, clonogenic content) using both univariable and multivariable models to distinguish between gross (unadjusted) associations and adjusted, independent effects. For each set of models, we assessed sensitivity by repeating the analyses after filtering out patients which belonged to a group with fewer than 3 members. Inclusion membership for the tests is provided in Additional file 1. For univariable analysis, Age was evaluated using Spearman’s correlation. Categorical variables were evaluated using Wilcoxon sum-rank for 2 groups and Kruskal-Wallis for 3 or more groups. For multivariable models, the data were fit to a linear model as described above and the contributions of each variable were evaluated using a Type-II ANCOVA on the linear model.

| **Robustness Level** | **Criteria** |
| --- | --- |
| **Not robust** | No significant association or inconsistent across models. |
| **Low** | (1) P-value < 0.05 in at least one univariable model<br>OR<br>(2) P-value < 0.2 for every test |
| **Moderate** | (1) P-value < 0.05 in at least one multivariable model<br>OR<br>(2) P-value < 0.2 in multivariate AND inclusion in low evidence robustness group 1 |
| **High** | (1) P-value < 0.05 in at least one multivariable model AND inclusion in at least one other low or moderate evidence robustness group |

#### Proteomic data modeling

Imputed, intra-MEC data matrices were analyzed independently. Estimates were calculated using a linear model with Age as a continuous variable and the remaining variables (Genotype, Parity, Phase and subsequently Colony Count) as categorical variable. GSEA was applied using a preranked list of Gene Symbols sorted based on estimated fold changes against the Gene Ontology Biological Processes, KEGG Pathways, and Cancer Hallmark gene sets with minimum and maximum sizes of 25 and 200, respectively. Outputs from GSEA were used as inputs to Cytoscape (v.3.9.1). GSEA gene sets were visualized in EnrichmentMap (v.3.3.4) and AutoAnnotate (v.1.3.5).

For TCGA breast cancer samples, the RNA Seq classifications of PAM50 subtype were used, except for those re-classified as Claudin-low in a re-analysis [34].

### Data visualization

Unless otherwise specified, plots were generated using R programming language (v.4.2.2) with Tidyverse package (tidyverse_1.3.2) with ggthemes_4.2.4, ggpubr_0.5.0, ggplot2_3.4.0, and ggbeeswarm_0.6.0 packages.

## Acknowledgements

Work in the Kislinger and Khokha labs was supported by operating grants from the Canadian Institutes of Health Research, the Terry Fox Research Institute PPG, Canadian Cancer Society, the Canadian Research Chair program. M.W. was supported by a CIHR Postdoctoral Fellowship. We would like to thank Miquel Angel Pujana for the careful review of the manuscript and Curtis W. McCloskey for his organization of the cohort data. Flow cytometry was performed using instrumentation in the Princess Margaret Flow facility.

## Author Contributions

Ma.W., B.Z., P.T., M.G, A.K., Mi.W., and P.W. designed experiments, performed experiments, and contributed to data acquisition. Ma.W. and Mi.W. acquired the mass spectrometry data. M.W. and B.Z. analyzed the data. Ma.W. generated figures. R.K. and T.K. supervised the study. Ma.W. and B.Z. wrote the manuscript, which all other authors edited and approved.

## Competing interests

Authors have no competing interests to declare.

## Additional information

Supplementary Information is available for this paper. Correspondence and requests for materials should be directed to and will be fulfilled by lead contacts, Rama Khokha and Thomas Kislinger.

## Data availability

Processed proteomics data are available in this paper’s Additional file 2. Any additional information required to reanalyze the data reported in this paper is available from the lead contact upon request.

---

**Figure S1.**
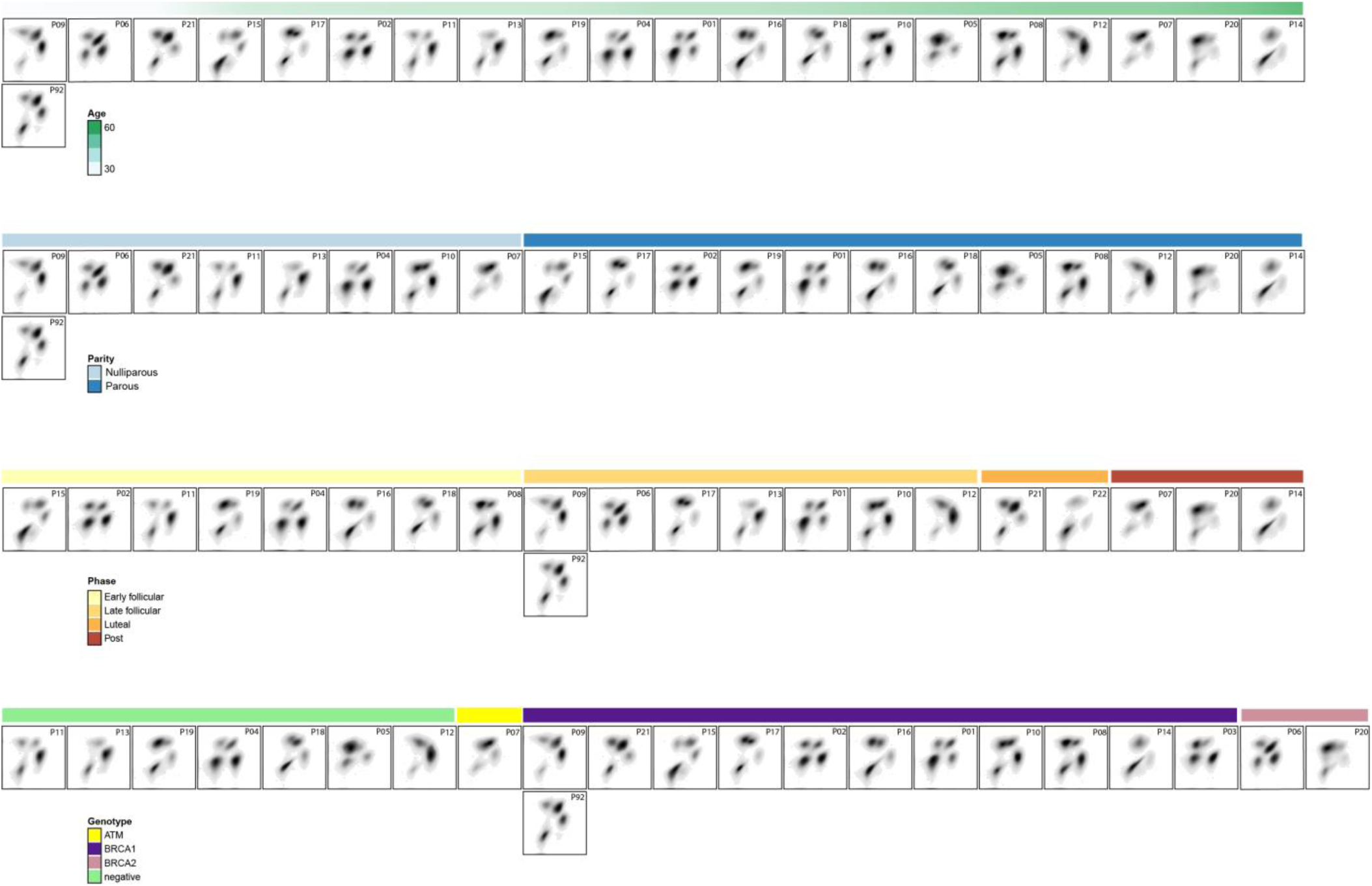
Donor flow cytometry profiles reveal inter-individual variability in MEC subtype separation. Flow cytometry density plots for all donors, ordered by clinical covariates. A possible relationship between donor age and degree of MEC separation is visible, though not statistically resolved.

**Figure S2.**
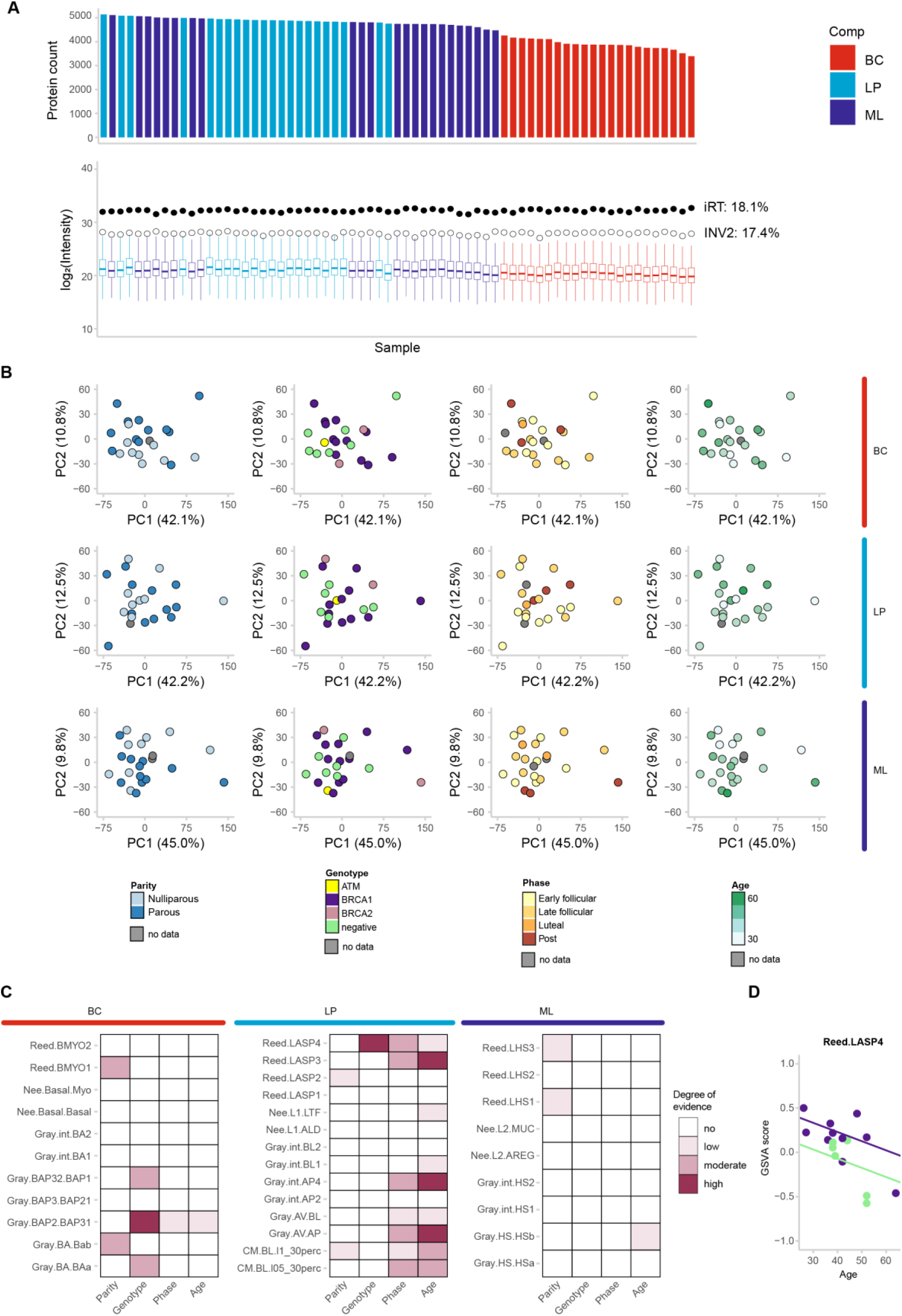
Clinical covariates influence enrichment of previously reported MEC subpopulation signatures. (A) Technical reproducibility and proteomic depth across runs. (B) Subtype-specific PCA for BC, LP, and ML cells. (C) Evidence scores for associations between subpopulation signatures (Reed, Nguyen, Pal) and clinical covariates. (D) Distribution of LASP4 enrichment scores plotted against age and germline mutation status.

**Figure S3.**
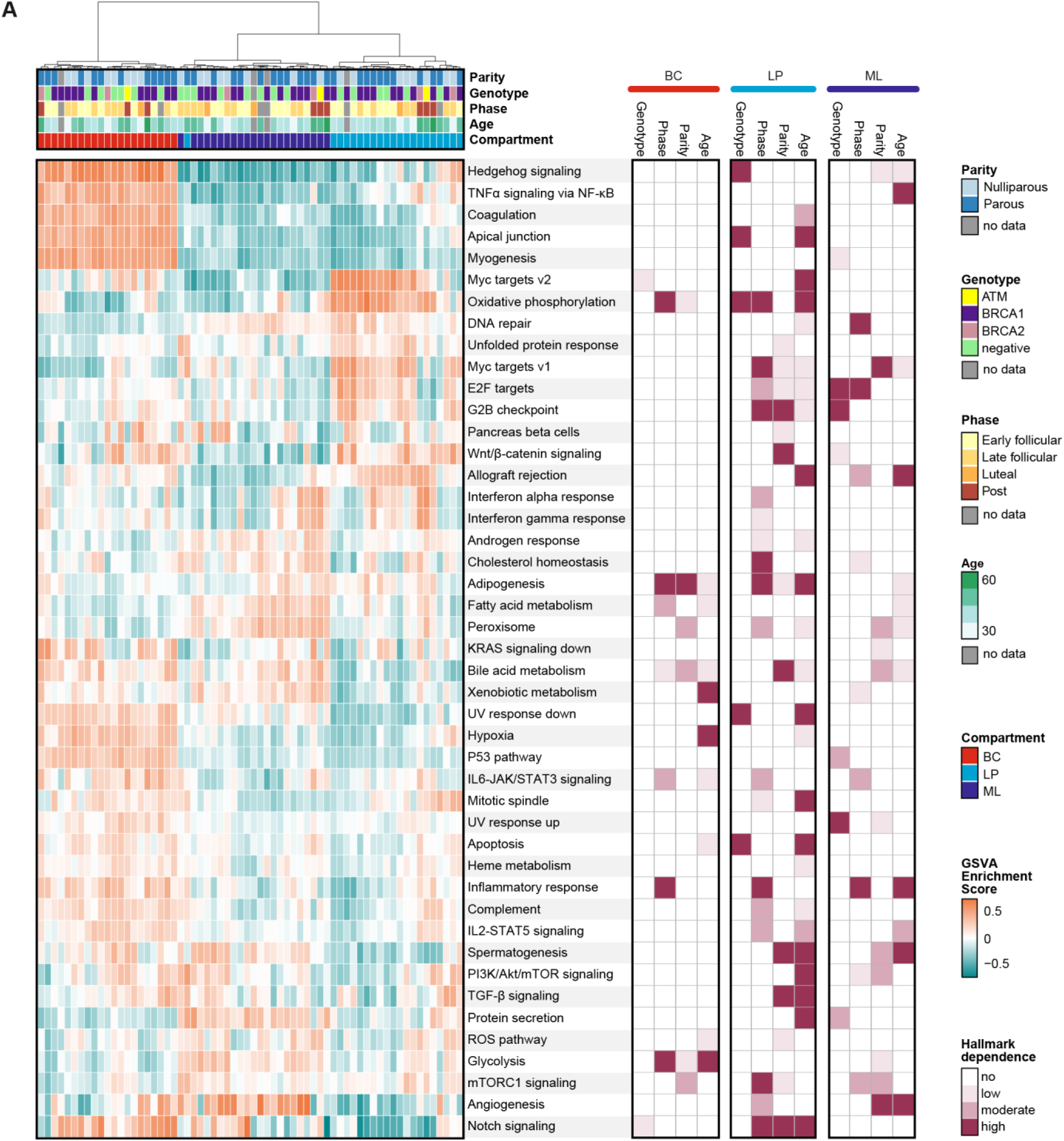
Sample-level Hallmark Signature enrichment highlights subtype-specific pathway programs and clinical covariate associations. Heatmap and clustering of Hallmark GSVA scores across donors for each MEC subtype. Level of evidence for association with clinical covariates is plotted.

**Figure S4.**
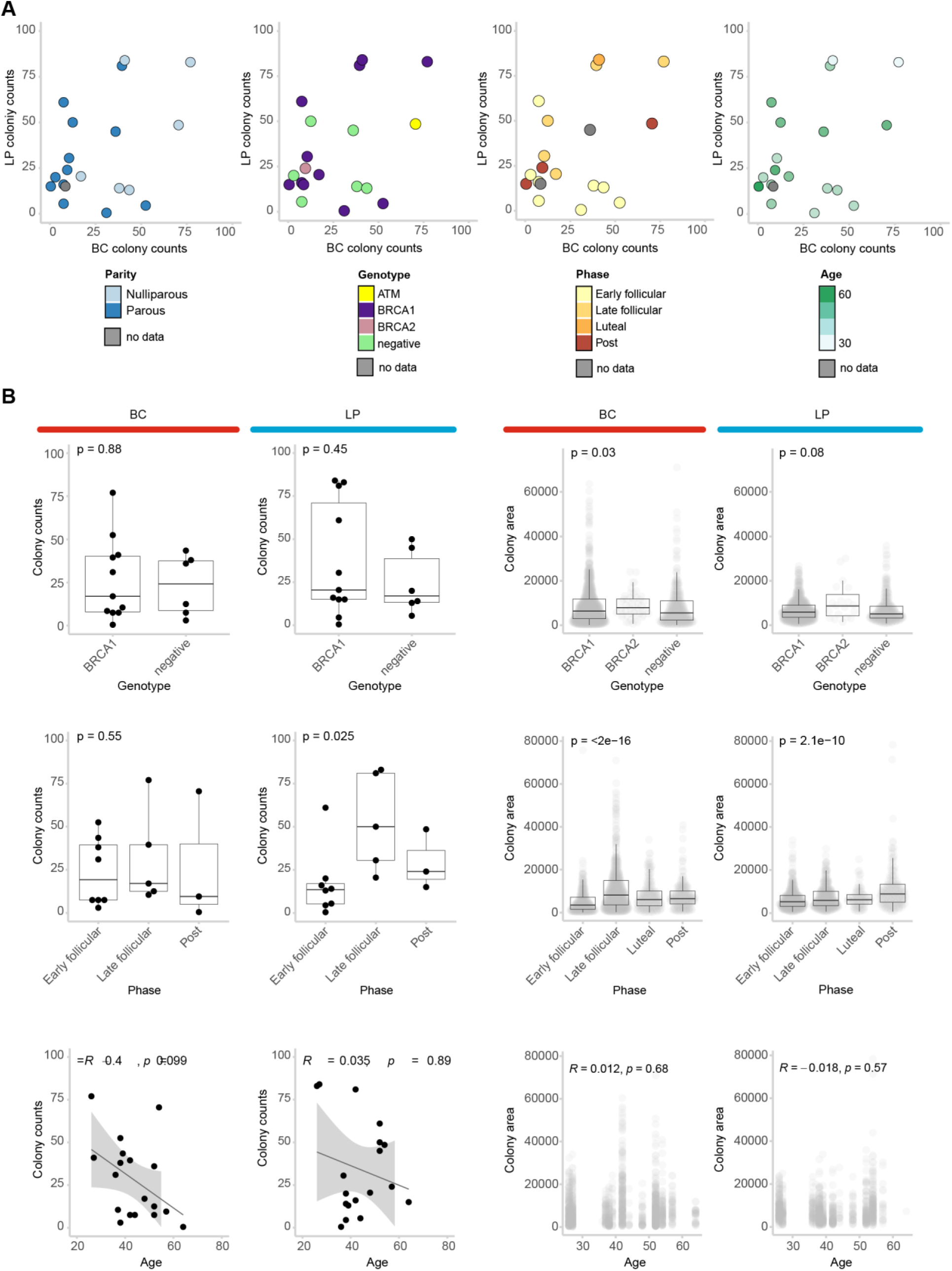
Assessing how clinical covariates impact clonogenicity. (A) Plots of BC and LP colony counts annotated with clinical covariates. (B) Distributions of colony counts and colony sizes across clinical covariates with univariable analysis results. Wicoxon rank sum test is used for comparison of two categories, Kruskal-Wallis is used for comparison of three or more categories. Unadjusted p-values are displayed for each test. Boxplots show median and interquartile values, and whiskers show 95% confidence intervals estimates. Spearman’s correlation R and p-values are displayed with a linear model estimate where shading indicated 95% confidence interval. Summary of additional statistical tests are provided in Additional file 1.

## Additional files

Additional file 1: Statistic summaries for univariate and multivariate models, related to all Figures

Additional file 2: Proteomics results table for high-risk breast cohort, related to Figure 2-4

Additional file 3: Complete GSEA results, related to Figure 3, 4

## Notes

### Competing Interest Statement

The authors have declared no competing interest.

